# Discrimination of Jamaican fruit bat lymphocytes by flow cytometry

**DOI:** 10.1101/2024.07.18.604131

**Authors:** Bradly E. Burke, Savannah M. Rocha, Corey Campbell, Elizabeth Creissen, Ronald B. Tjalkens, Wenjun Ma, Marcela Henao-Tamayo, Tony Schountz

**Affiliations:** Department of Microbiology, Immunology and Pathology, College of Veterinary Medicine and Biomedical Sciences, Colorado State University, Fort Collins, CO, 80523 USA; Flow Cytometry and Cell Sorting Core Facility, College of Veterinary Medicine and Biomedical Sciences, Colorado State University, Fort Collins, CO, 80523 USA; Department of Environmental and Radiological Health Sciences, College of Veterinary Medicine and Biomedical Sciences, Colorado State University, Fort Collins, CO, 80523 USA; Department of Veterinary Pathobiology, College of Veterinary Medicine and Department of Molecular Microbiology and Immunology, School of Medicine, University of Missouri, Columbia MO 65211 USA

## Abstract

Bats are natural reservoir hosts of many important zoonotic viruses but because there are few immunological reagents and breeding colonies available for infectious disease research, little is known about their immune responses to infection. We established a breeding colony Jamaican fruit bats (*Artibeus jamaicensis*) to study bat virology and immunology. The species is used as a natural reservoir model for H18N11 influenza A virus, and as a surrogate model for SARS-CoV-2, MERS-CoV and Tacaribe virus. As part of our ongoing efforts to develop this model organism, we sought to identify commercially available monoclonal antibodies (mAb) for profiling Jamaican fruit bat lymphocytes. We identified several cross-reactive mAb that can be used to identify T and B cells; however, we were unable to identify mAb for three informative T cell markers, CD3γ, CD4 and CD8α. We targeted these markers for the generation of hybridomas, and identified several clones to each that can be used with flow cytometry and fluorescence microscopy. Specificity of the monoclonal antibodies was validated by sorting lymphocytes, followed by PCR identification of confirmatory transcripts. Spleens of Jamaican fruit bats possess about half the number of T cells than do human or mouse spleens, and we identified an unusual population of cells that expressed the B cell marker CD19 and the T cell marker CD3. The availability of these monoclonal antibodies will permit a more thorough examination of adaptive immune responses in Jamaican fruit bats that should help clarify how the bats control viral infections and without disease.

**Importance:** Bats naturally host a number of viruses without disease, but which can cause significant disease in humans. Virtually nothing is known about adaptive immune responses in bats because of a lack of immunological tools to examine such responses. We have begun to address this deficiency by identifying several commercially available monoclonal antibodies to human and mouse antigens that are cross-reactive to Jamaican fruit bat lymphocyte orthologs. We also generated monoclonal antibodies to Jamaican fruit bat CD3γ, CD4 and CD8α that are suitable for identifying T cell subsets by flow cytometry and immunofluorescent staining of fixed tissues. Together, these reagents will allow a more detailed examination of lymphocyte populations in Jamaican fruit bats.

## Introduction

There are more than 1,400 species of bats that serve important ecological roles, including pollination and seed dispersal, and consumption of insects such as crop and forest pests, and mosquitos and midges that spread arthropod-borne diseases [1]. Several bat species are reservoir hosts of viruses that cause disease in humans and livestock, including, henipaviruses, filoviruses, coronaviruses and lyssaviruses [2]. SARS-CoV-2, the cause of the COVID-19 pandemic, likely originated from a horseshoe bat reservoir [3]. Moreover, bats are susceptible to disease from other pathogens such as rabies virus and the fungus *Pseudogymnoascus destructans* that causes white nose syndrome (WNS) that has killed millions of North American bats, and reduced three species (*Myotis septentrionalis*, *Myotis lucifugus*, and *Perimyotis subflavus*) by more than 90% [4].

Human and rodent immunology provide a foundation to understand immune systems and responses of other mammals; however, clear differences exist between these well understood species and bats. Some bat species have an “always on” innate type I interferon (IFN) system with higher basal expression of type I IFN, unlike humans and mice [5]. However, constitutive expression of IFNα in some bat species induces the expression of a subset of IFN-stimulated genes but without chronic inflammatory pathology [6]. In some bat species, the number of VDJ immunoglobulin heavy chain segments are expanded [7, 8] compared to humans, which have about 40 functional V segments, 24 D segments, and 6 J segments that can lead to more than 5,000 specificities through recombination alone [9]. The little brown bat (*Myotis lucifugus*), in contrast, has at least 236 V segments, 24 D segments, and 13 J segments giving rise to more than 73,000 heavy chain combinations [7]. Furthermore, little brown bats appear use affinity maturation of antibodies to a lesser degree than humans [7]. The Egyptian fruit bat (*Rousettus aegyptiacus*) has a similar repertoire potential as humans but its genome has an expanded KLRC/KLRD family of natural killer (NK) cell receptors, MHC class I genes, and type I interferons [10].

Despite the differences known about the immune system of bats, there is little known about the cellular immune systems of the different species and subsequent immune responses to infectious agents. Jamaican fruit bats (*Artibeus jamaicensis*) are native to the Caribbean and Central and South America and represent an underdeveloped animal model for immunology research. Jamaican fruit bats are a natural reservoir of H18N11 influenza A virus, and the species is used to study other viral infections as a surrogate host, including SARS-CoV-2, MERS-CoV, Zika virus, rabies virus and Tacaribe virus [11–15]. The lack of immunological reagents for bats constrains the use of high throughput technologies, such as flow cytometry and cell sorting, and quantitative data and anatomical determination from fluorescence microscopy. To address this gap, we first sought to identify commercially available monoclonal antibodies (mAb) that were cross-reactive with Jamaican fruit bat orthologs by *in silico* pre-screening protein homology for potential cross-reactivity, followed by flow cytometry and florescent microscopy. We then developed three mAb for which commercially available cross-reactive antibodies could not be identified. We validated specificity of target cells by flow cytometric sorting and performing quantitative reverse transcription PCR (qRT-PCR) for the target sequences. Together, we identified or generated more than a dozen antibodies that are useful for flow cytometry and immunofluorescence use with Jamaican fruit bats.

## Results

### Pre-screening antibody cross-reactivity

Jamaican fruit bat CD3ε protein homology from BLOSUM62 analysis indicated 71.63% homology with human CD3ε (**Figure 1A**) Phyre2 protein modeling of human and Jamaican fruit bat CD3ε and a second BLOSUM62 analysis between the two proteins indicated 75.5% homology with 133 (63.9%) identical sites (**Figure S1**). Jamaican fruit bat CD19 protein homology from BLOSUM62 analysis indicated 72% homology with mouse CD19 across all isoforms (**Figure 1A**). Phyre2 protein modeling of CD19 mouse and Jamaican fruit bat CD19 with a second BLOSUM62 analysis between the two proteins indicated 67.1% homology with 336 (56.9%) identical sites (**Figure S2**). Phyre2 modeling of Jamaican fruit bat CD3γ and CD4 had the least identity, suggesting mAb to other species are less likely to be cross-reactive (**Figure S3**). Unrooted tree of CD19, CD3γ and CD3ε cDNA sequences from **Figure 1A** showed clustering of New World bat sequences and clustering of Old World bat sequences (**Figure 1B**).

**Figure 1.**
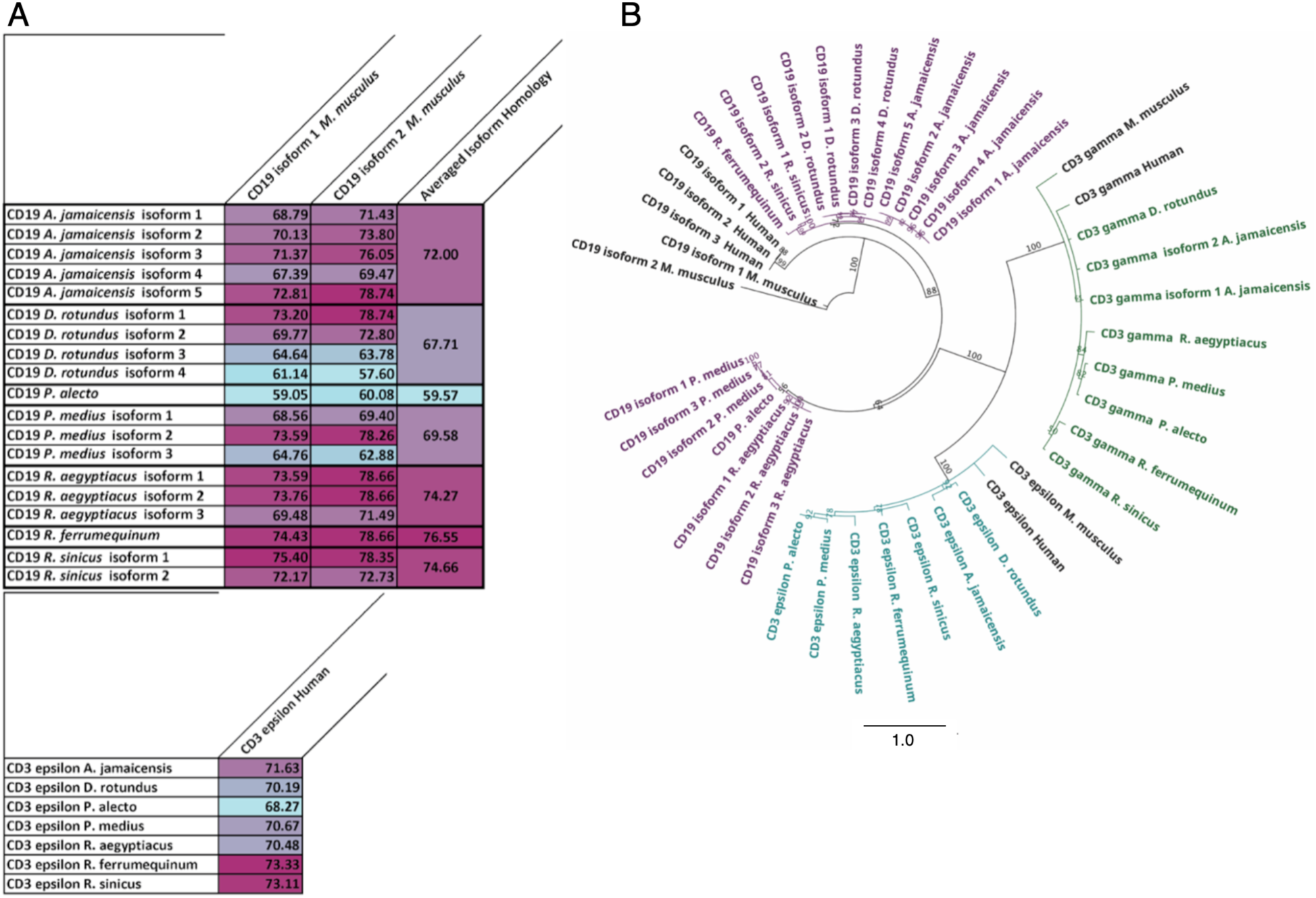
Protein sequence homology of CD19 and CD3ε isoforms. (A) Heat map of Geneious Protein Alignment BLOSUM62 analysis of human (*Homo sapiens*), mouse (*Mus musculus*), Jamaican fruit bat (*Artibeus jamaicensis*), common vampire bat (*Desmodus rotundus*), Chinese horseshoe bat (*Rhinolophus sinicus*), greater horseshoe bat (*Rhinolophus ferrumequinum*), Egyptian fruit bat ( *Rousettus aegyptiacus*), Indian flying fox (*Pteropus giganteus*), and black flying fox (*Pteropus alecto*) for all CD19 and CD3e protein isoforms [29, 30]. Cyan indicates lowest homology percentage, and plum indicates highest homology percentage. (B) Unrooted tree generated from Geneious Protein Alignment BLOSUM62 analysis of human (*Homo sapiens*), mouse (*Mus musculus*), Jamaican fruit bat (*Artibeus jamaicensis*), common vampire bat (*Desmodus rotundus*), Chinese horseshoe bat (*Rhinolophus sinicus*), greater horseshoe bat (*Rhinolophus ferrumequinum*), Egyptian fruit bat ( *Rousettus aegyptiacus*), Indian flying fox (*Pteropus giganteus*), and black flying fox (*Pteropus alecto*) for all CD3ε, CD3γ, and CD19 protein isoforms [29, 30]. Branch numbers indicated bootstrap percentages.

### Immunofluorescence validation of antibody reactivity and anatomical mapping of immune cells

Cross-reactivity of CD19 1D3, CD3ε Hit3a, and CD3γ XE-2 were validated through immunofluorescent staining within the small intestine. Anatomical localization of cellular populations was determined through 20x magnification montage imaging of whole tissue sections (**Figure 2A, 2E, 2I**). Positive identification of antigen specific staining was performed through peri-nuclear localization of CD3γ (**Figure 2B, 2K**), CD3ε (**Figure 2C, 2G**), and CD19 (**Figure 2F, 2J**) with DAPI^+^ staining. Staining colocalization of CD3γ^+^ and CD3ε^+^ (**Figure 2B-2D**) and CD19^+^ and CD3ε^+^ (**Figure 2F-2H**) cells were observed in the lamina propria of the intestinal villi (**Figure 2A, 2E**). Cells containing colocalization of CD19^+^ and CD3γ^+^ staining (**Figure 2J-2L**) were observed within evenly spaced, condensed cellular formations located below intestinal crypts. In relation to intestinal lumen, these cellular aggregates are superficial to the submucosal, inner muscle and outer muscle sublayers, therefore, resembling that of lymphatic nodules (micro-Peyer’s patches, **Figure 2I**).

**Figure 2.**
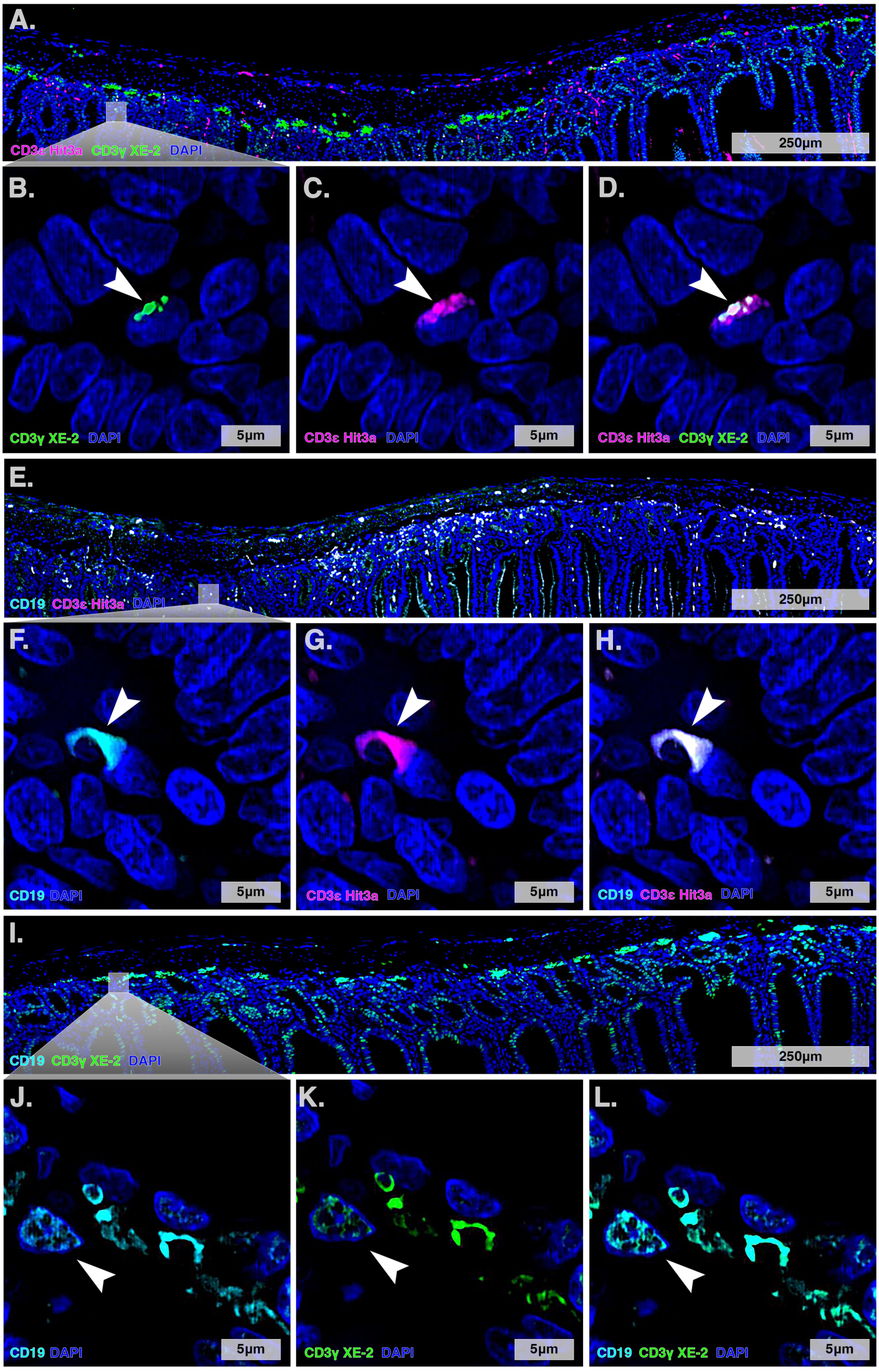
Immunofluorescence identification and anatomical localization of B and T-cell populations in the small intestine of Jamaican fruit bats. Low magnification (20X) montage imaging of the small intestine in male Jamaican fruit bats highlighting the anatomical location of CD3ε^+^ CD3γ^+^ cells (A), CD19^+^ CD3ε^+^ cells (**E**), and CD19^+^ CD3γ^+^ cells (**I**). High magnification (100X) Z-stack images with applied deconvolution and projection parameters show individual channel positivity for the staining combinations and arrowheads indicate cells positive for the respective dual staining parameters (**B-D, F-H, J-L**). Colocalization of CD3γ^+^ CD3ε^+^ (**D**) T-cells with respective individual channel representations (**B, C**). Colocalization of rare CD19^+^ CD3ε^+^ cells within the lamina propria of a transverse section of the intestinal villi (**H**) with respective individual channel representations (**F, G**). Colocalization of a second rare population of CD19^+^ CD3γ^+^ cells (**L**) with respective individual channel representations of CD19 (**J**) and CD3γ (**K**).

### Cell Sorting

Cells were enriched by sorting into four enriched categories by a gating strategy (**Figure 3**): Not sorted (whole spleen control), CD19^+^, CD3ε^+^, CD3γ^+^, and CD3ε^+^ CD3γ^+^. Cell counts determined by FACSDiva and RNA yielded by Trizol extraction and UV spectrometry for each enriched population (**Table 1)**. Minimum number of cells sorted was 2,543 and maximum number of cells sorted was 1,243,632. Minimum RNA yield was 245.20 ng and maximum RNA yield was 1,318.15 ng (**Table 1**).

**Figure 3.**
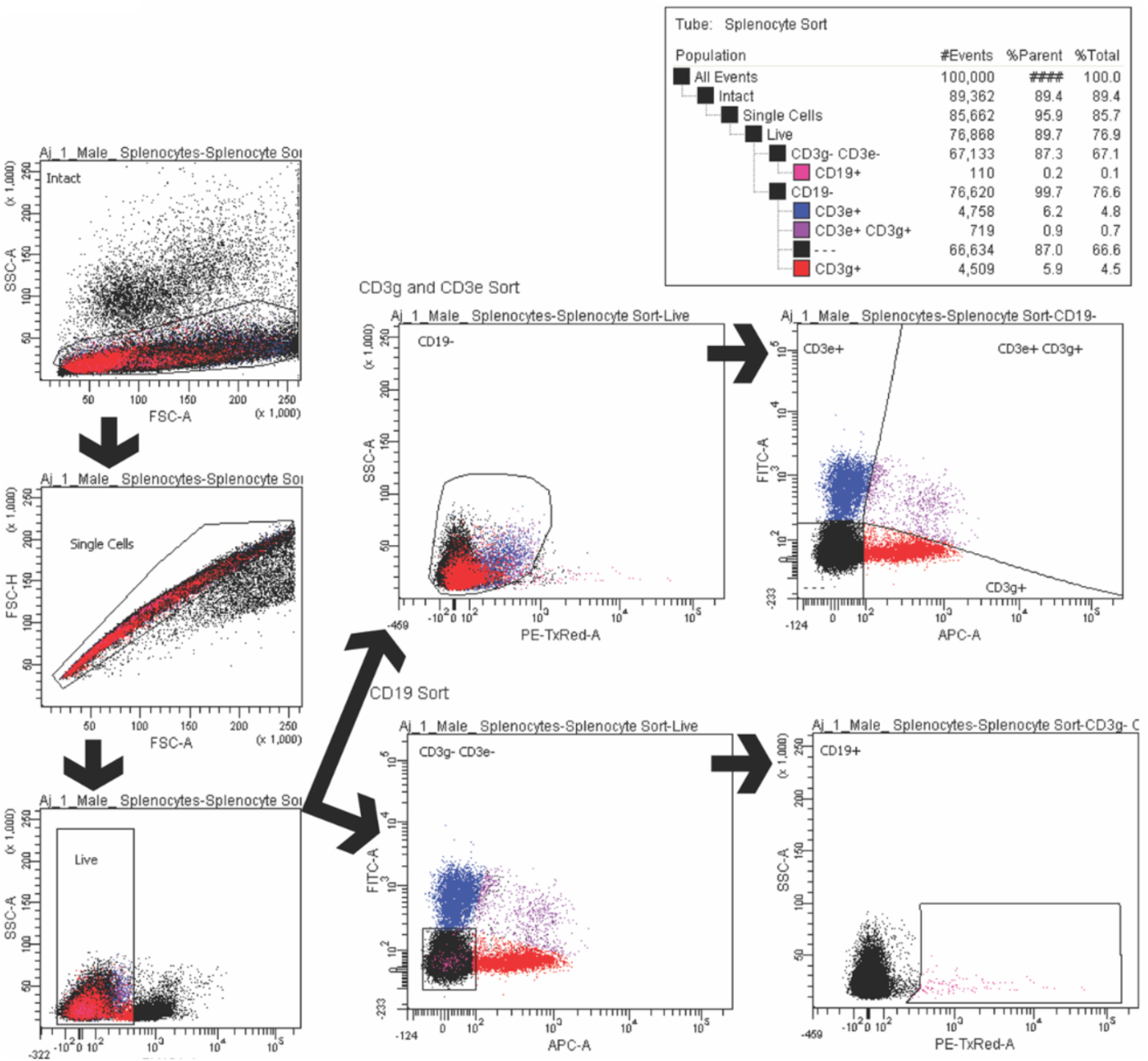
Cell sorting Jamaican fruit bat splenocytes. Splenocytes were then stained with Tonbo Ghost Violet 450 viability dye followed by staining with CD3γ (Clone: XE2, Fluorophore: Alexa Fluor 647), CD3ε (Hit3a, FITC), and CD19 (1D3, PE-eFluor610). Cells were sorted into six categories using a BD FACSAria-III: Not sorted (whole spleen), CD19^+^, CD3ε^+^, CD3γ^+^, and CD3ε^+^ CD3γ^+^. Gating strategy used initial clean-up gates: Intact lymphocytes > Single Cells > Live Cells followed by CD19^-^ or CD3ε^-^ CD3γ^-^. CD19-cells were then sorted into CD3ε^+^, CD3γ^+^, or CD3ε^+^ CD3γ^+^ populations. CD3ε^-^ CD3γ^-^ cells were then sorted into a CD19^+^ population.

**Table 1.**
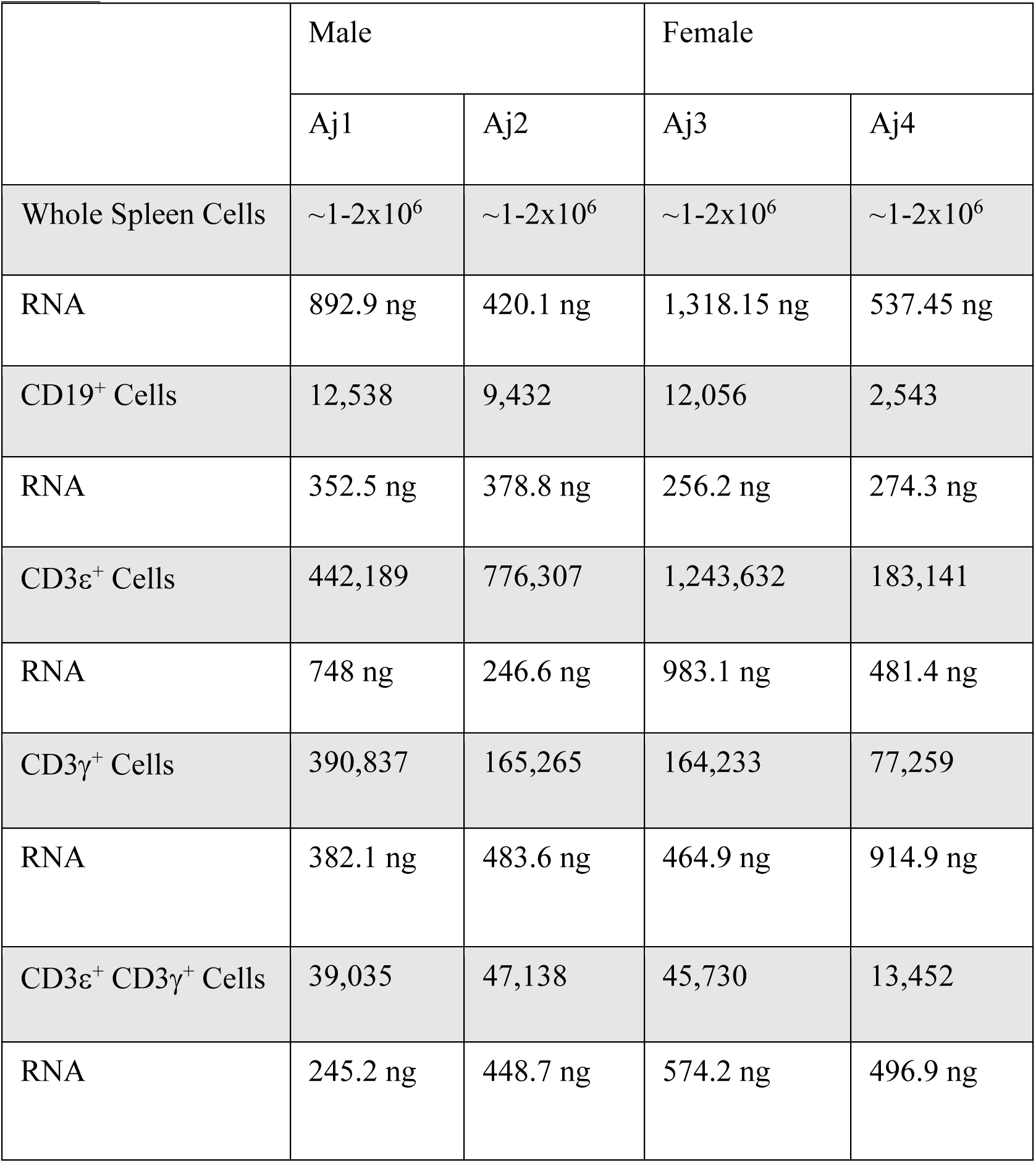

### Post-Hoc Cell Sorting Analysis

Concatenated data was gated two ways, either first gating on CD3ε^+^, CD3γ^+^, CD3ε^+^CD3γ^+^ followed by CD19 gating, or first gating on CD19^+^ cells followed by CD3ε^+^, CD3γ^+^, and CD3ε^+^CD3γ^+^ gating (**Figure 4**). This demonstrated that CD3ε^+^ cells had minimal CD19^+^ cells 0.073%. CD3ε^+^CD3γ^+^ cells had 1% CD19^+^ cells. CD3γ^+^ cells had 17% CD19^+^ cells. CD19^+^ cells contained 2.23% CD3ε^+^ cells, 3.79% CD3ε^+^CD3γ^+^ cells, 30.1% CD3γ^+^ cells, and 63.8% CD3ε^-^ CD3γ^-^ cells.

**Figure 4.**
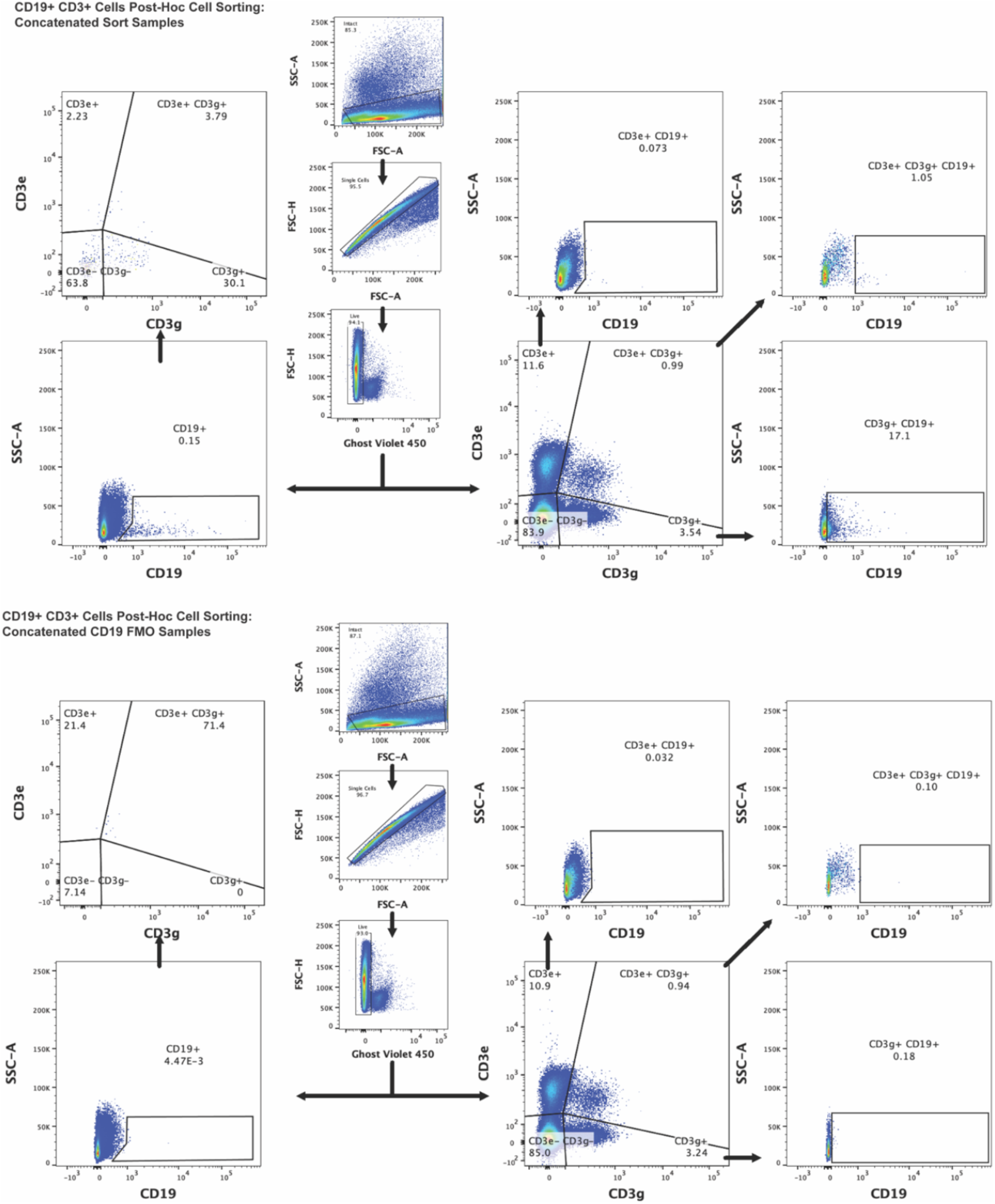
Concatenated post-hoc cell sorting analysis. FCS files generated from cell sorting were analyzed in FlowJo (v10.8.1). FCS files were from all four bats were concatenated to better visualize CD19^+^ CD3^+^ cell populations. The top portion of the figure shows concatenated sorted populations, and the bottom portion shows concatenated CD19 FMO samples. Gates were drawn using FMOs. Initial gating is as follows: Intact > Single Cells > Live. CD19^+^ CD3^+^ cells were then viewed two ways: 1) first gating on CD3^+^ cells followed by CD19^+^ cells 2) first gating on CD19^+^ cells followed by CD3^+^ cells.

### Validation of Antibodies by RT-qPCR

CD3ε^+^ splenocytes indicated significant gene expression increase for *Cd3ε* (♀), *Cd3γ* (♀), *Cd28* (♀) and *Cd45* (♀) (**Figures 5A, 5B, 5F, 5G**). CD3ε^+^ splenocytes had a significant decrease in *Cd19* (pooled, ♂ and ♀) (**Figure 5E**). CD3γ^+^ splenocytes had a significant gene expression increase for *Cd3ε* (pooled and ♂), *Cd3γ* (♀), *Cd8a* (♀), *CD28* (pooled, ♂, and ♀), and *Cd45* (♀) (**Figure s5A, 5B, 5F, 5G**). CD3ε^+^ CD3γ^+^ splenocytes had significant gene expression increases for *Cd3ε* (pooled, ♂, ♀), *Cd3γ* (pooled, ♂, ♀), *Cd4* (pooled, ♂, ♀), *Cd8a* (pooled, ♂, ♀), *Cd28* (pooled, ♂, ♀), and *Cd45* (pooled and ♀) (**Figures 5A, 5B, 5C, 5D, 5F, 5G**). CD19^+^ splenocytes had significant gene expression increases for *Cd3γ* (♀), *Cd19* (pooled, ♀), and *Cd28* (pooled, ♂, ♀) (**Figures 5B, 5E, 5F**).

**Figure 5.**
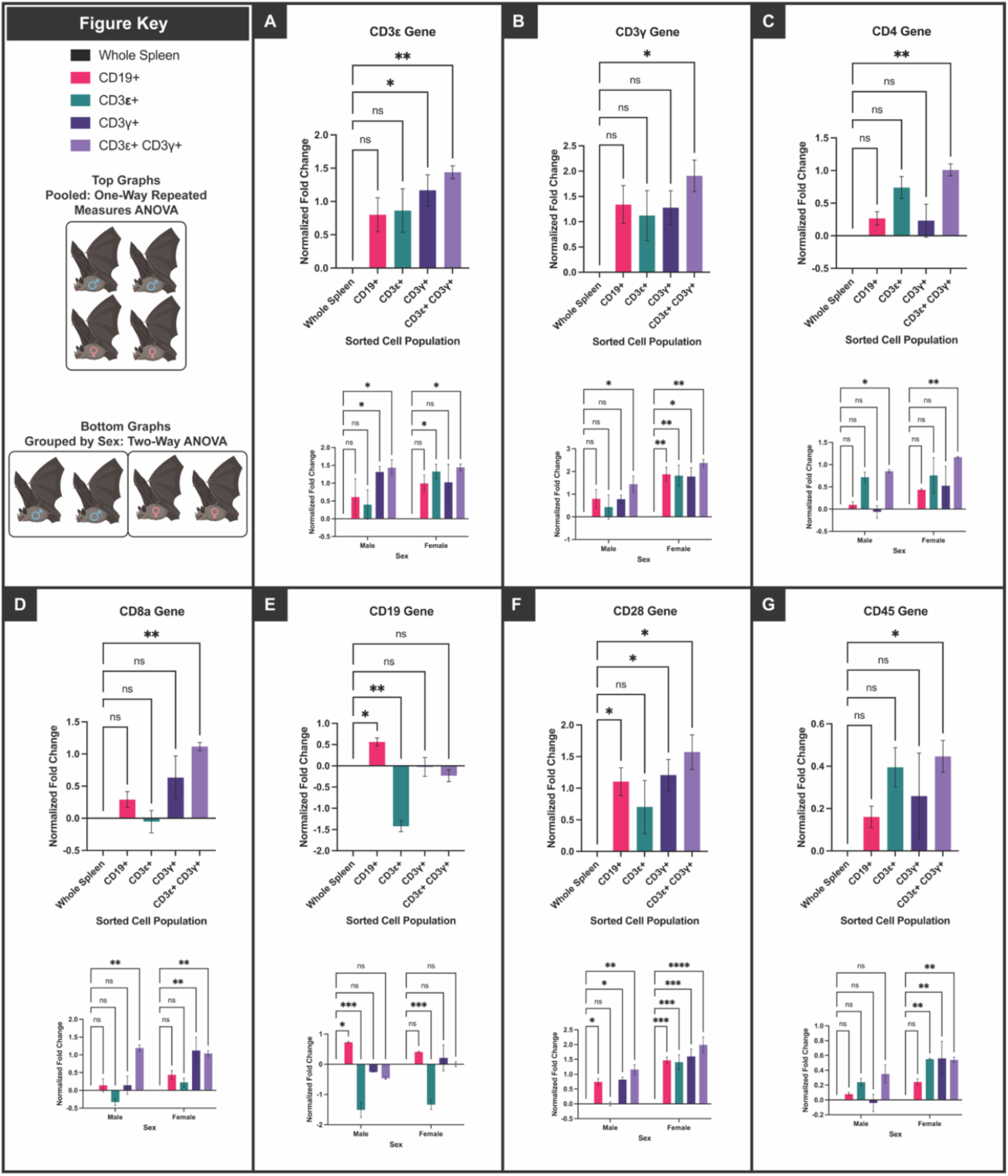
RT-qPCR analysis of not sorted (whole spleen), CD19^+^, CD3ε^+^, CD3γ^+^, and CD3ε^+^ CD3γ^+^ Jamaican fruit bat splenocytes. CD Normalized fold change to whole spleen control of RT-qPCR Data. For each panel, the top graph represents the pooled data out-put of the four Jamaican fruit bats used 2 males and 2 females. One-way repeated measure ANOVA was used for statistical analysis. The bottom graph represents the data output of the four Jamaican fruit bats grouped by sex, 2 males and 2 females. (**A**) *CD3*ε gene, (**B**) *Cd3γ* gene, (**C**) *Cd4* gene, (**D**) *Cd8a* gene, (**E**) *Cd19* gene, (**F**) *Cd28* gene, (**G**) *Cd45* gene.

### Generation of monoclonal antibodies to Jamaican fruit bat CD4, and CD8α

Hybridoma production targeting Jamaican fruit bat CD4 and CD8α peptide conjugates resulted in several dozen reactive clones for each antigen. CD4 clone 1-D5 showed 15.7% of cells expressed CD4 whereas clone 1-B2 showed 8.59% with varying reactivity with lymphocytes and monocytes based on forward scatter area light (FSC-A) by side scatter light area (SSC-A) (**Figures 6A and 6B**). Monocytes are larger (FSC-A) and have more granularity (SSC-A) than lymphocytes but less than granulocytes. Splenocytes stained with CD8α clone 2-H9 showed 7.39% positive and clone 2-H8 showed 11.6% positive (**Figures 6C and 6D**). Both CD8α clones when used to stain splenocytes the FSC-A and SSC-A profiles suggest predominantly lymphocytes were stained.

**Figure 6.**
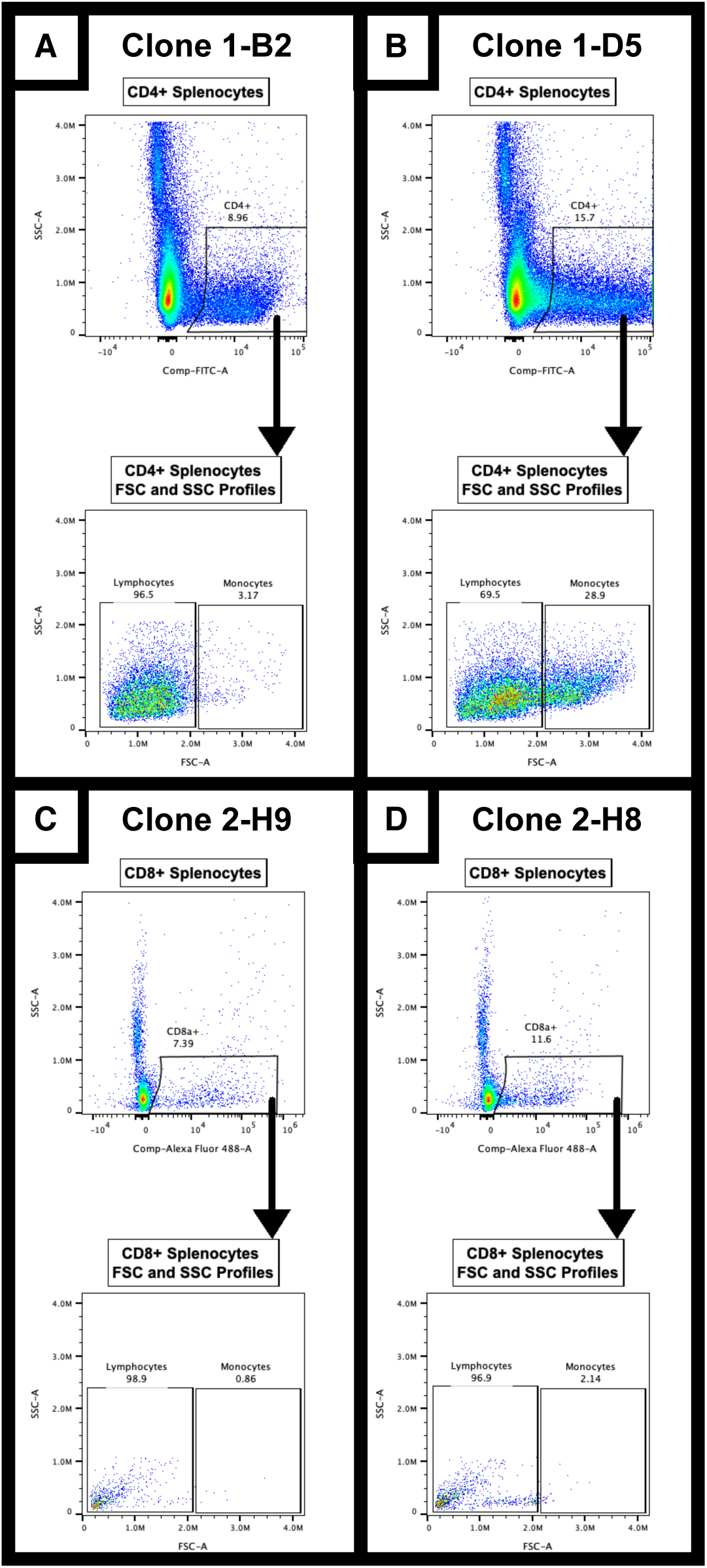
Variation of CD4 hybridoma clone reactivity with Jamaican fruit bat splenocytes. (A) Clone 1-D5 had high reactivity to both lymphocytes monocytes, whereas (B) clone 1-B2 had high reactivity predominantly with lymphocytes. Gating strategy: Intact cells > Single cells > Live cells > CD4^+^ cells> Lymphocytes / Monocytes.

## Discussion

The lack of adequate immunological reagents available to identify immune cells in bats limits the ability of investigation into their immune systems. This impairs the study of immune responses to bat viruses that cause substantial disease burden in humans (e.g. SARS-CoV-2, Marburg virus, Nipah virus). To address some of these challenges, we established a closed breeding colony of Jamaican fruit bats nearly twenty years ago for use in the study of bat-borne viral infections. Although many studies of wild and captive bats have shed light on the virology of infection, few have examined bat immune responses to viruses. Of those studies that did, nearly all examined the innate immune response using bat cells infected with viruses or sera to assess antibody responses. To date, investigations into the adaptive immune systems of bats have not been thoroughly conducted, principally due to the lack of validated reagents. We have identified and developed such reagents that will permit detailed examination of bat adaptive immune responses to viruses.

In CD3ε^+^ cells, a trending increase of *Cd3ε*, *Cd3γ*, *Cd4*, *Cd28*, and *Cd45* gene expression was observed, consistent with helper T cell phenotypes (**Figures 5A, 5B, 5C, 5F, 5G**). Of interest, CD3ε^+^ cells trended with a higher *Cd4* gene expression and a lower *Cd8a* gene expression (**Figures 5C and 5D**). This trend suggested that CD3ε ^+^ cells of Jamaican fruit bats largely contain CD4^+^ T cells, or that CD4 expression is higher in cells than is CD8 expression. In CD3γ^+^ cells, a trending increase of *Cd3γ*, *Cd8a*, and *Cd45* gene expression was observed, consistent with what would be expected of T cells (**Figures 5B, 5D, 5G**). In contrast to CD3ε^+^ cells, CD3γ^+^ cells trended with a higher *Cd8a* gene expression and a lower *Cd4* gene expression **(Figures 5C and 5D)**. This trend suggested that CD3γ^+^ cells of Jamaican fruit bats predominantly contained CD8α^+^ T cells. For the CD3ε^+^ CD3γ^+^ splenocyte population, gene expression was increased for *Cd3ε* (pooled, ♂, ♀), *Cd3γ* (pooled, ♂, ♀), *Cd4* (pooled, ♂, ♀), *Cd8a* (pooled, ♂, ♀), *Cd28* (pooled, ♂, ♀), and *Cd45* (pooled and ♀). CD3ε^+^ CD3γ^+^ splenocyte population of Jamaican fruit bat T cells suggested the presence of canonical CD4 and CD8 T cell populations (**Figures 5A, 5B, 5C, 5D, 5F, 5G**). Normal human spleens contain 31% CD3ε^+^ cells [16], whereas normal mouse spleens contain roughly 30% CD3ε^+^ cells [17–19]. CD3ε is commonly used for T lymphocyte lineage due to the establishment of the TCRαβ, CD3δɛ, CD3γɛ dimer stoichiometry 2(ɛ)/1(δ)/1(γ), such that CD3ɛ is the most abundant CD3 chain [20, 21]. Compared to human and mouse spleens, containing roughly 30%

CD3ε^+^ T lymphocytes, the four Jamaican fruit bats averaged 11.6% CD3ε^+^ T lymphocytes CD3γ^+^ T lymphocytes accounted for 3.54% of the concatenated Jamaican fruit bat spleens. CD3ε^+^, CD3γ^+^, and CD3ε^+^ CD3γ^+^ T lymphocytes averaged across the four Jamaican fruit bats, with 16.13% (**Figure 4**); half that is observed in human and mouse spleens. It has been shown using a cross-reactive intracellular anti-human CD3ε antibody, black flying fox spleens are roughly 15.6% CD3ε^+^, but no other CD3 chains were targeted for labeling [22]. As such, the established TCRαβ, CD3δɛ, CD3γɛ dimer stoichiometry 2(ɛ)/1(δ)/1(γ) in Jamaican fruit bats, and likely other bat species, is unknown. Furthermore, the implication of this study highlights the importance and persistent need for continued investigation into Jamaican fruit bat T lymphocyte abundance and why it is nearly half that of human and mouse spleens. This could be due to a unique CD3 dimer stoichiometry specific to these animals. However, it is possible that the Jamaican fruit bat T cell populations cannot be visualized with cell surface staining in its entirety without labeling CD3δ. Given this, the currently accepted assumption that bat, human and mouse dimer stoichiometry is equivalent may lead to erroneous populational representations of T cells and T cell subsets when investigating bat immune cell populations. The results herein reveal that identification of T cell populations by implementation of CD3ε staining, a recognized standard pan T cell marker, would only account for a mere subset of T cells, where ∼3.5% of CD3^+^ cells would be missed. Likewise, if identification of T cell populations were solely based upon anti-CD3γ, ∼11.6% of CD3^+^ cells would not be detected. As such, anti-CD3δ, anti-CD3ζ, anti-TCRα, and anti-TCRβ antibodies specific for detection in Jamaican fruit bats are needed to accurately and effectively characterize T lymphocyte and other CD3^+^ cell populations. In addition to the ambiguity of CD3 dimer stoichiometry in bats, it is noteworthy that CD3ε^+^ population trended towards a CD4 T helper compartment upon gene expression analysis (**Figure 5C**) and the CD3γ^+^ population trended towards a CD8 cytotoxic T cell compartment, especially in female bats (**Figure 5D**). As such, bat specific CD4 and CD8α monoclonal antibodies were generated to further explain how CD3γ^+^ and the CD3ε^+^ populations lean in respect to CD4 and CD8 protein expression in future studies (**Figure 5C and 5D**).

Analysis of CD19 splenocyte populations revealed significant gene expression increases for the following: *Cd3γ* (♀), *Cd19* (pooled, ♀), and *Cd28* (pooled, ♂, ♀) (**Figures 5B, 5E, 5F**). The data also indicated a trending increase of *Cd3ε* and *Cd3γ* gene expression. (**Figures 5A, 5B**) The cell sorting gating strategy attempted to remove CD3^+^ CD19^+^ cells by first gating on CD3ε^-^ CD3γ^-^ splenocytes (**Figure 3**). CD19^+^ CD3^+^ cells were also identified through immunofluorescence co-staining in the small intestine (**Figure 2**). In concert, three distinct biochemical and molecular detection-based assays (flow cytometry, fluorescence microscopy, and RT-qPCR) uncovered a novel subset of CD19^+^ CD3^+^ cells (**Figures 2, 4, 5**). The exact functionality/physiological role of this distinct cell type should be an important focus of future research. These may be a subset of T cells rather than a subset of B cells because they express CD3 proteins as shown by flow cytometry and immunofluorescence microscopy, in combination with statistically significant increased gene expression of *Cd28*, another T cell marker (**Figures 2, 4, and 5F**). In humans, a case of a novel CD19-expressing T-cell lymphoma has been documented in a 29 year old male [23]. Additionally, dual positive CD19^+^ CD3^+^ CAR T-cells have also been identified [24]. As such, there is evidence supporting that CD19 expression can occur on certain T cells, perhaps rarely in humans or mice, even though CD19 expression on T cells is not typical of a homeostatic immune state of humans or mice. Additionally, bats are a long-lived species and have low incidences of cancer [25, 26]. As such, this suggests that this novel subset of CD19^+^ CD3^+^ cells are a part of the Jamaican fruit bat homeostatic immune system. CD19 interacts with the complement system by dimerizing with CD21 complement C3 receptor and lowers the activation threshold on B cells [27, 28]. As such, a possible function for this subset of CD19^+^ CD3^+^ T cells may be activation by opsonized antigens using the complement system, or similarly lowers the activation threshold as in B cells.

Furthermore, with antibody clones 1D3 anti-CD19 and Hit3a anti-CD3ε demonstrating cross-reactivity with Jamaican fruit bat epitopes it is likely that these clones will cross-react with other bat species as indicated by BLOSUM62 protein homology (**Figure 1**). This work will help provide a framework for other researchers in the field of bat immunology to more rigorously test antibody cross-reactivity and help build a more robust body of literature of cross-reactive antibodies.

## Conclusions and Future Directions

The paucity of bat specific reagents makes cross-reactive validation studies vital to discover commercially available reagents that can be utilized and/or the lack thereof. This study has demonstrated two commercially available anti-mouse antibodies, CD3ε (clone Hit3a) and CD19 (clone 1D3), that cross-react with Jamaican fruit bat orthologs. Validation of these antibodies in fresh and fixed tissue allows for their use in flow cytometry assays and fluorescent microscopy. Validation of these reagents gives additional tools to help investigators elucidate the differences in the Jamaican fruit bat homeostatic immune system and immune responses compared to established animal models and clinical research conducted in humans. However, it is to be noted and warrants further study as to why Jamaican fruit bat splenic T lymphocyte populations (as well as black flying foxes) are nearly half that of humans and mice. Furthermore, this study highlights the need to develop more Jamaican fruit bat specific reagents because it appears that Jamaican fruit bat CD3 dimer stoichiometry might not follow the canonical ratios of humans and mice. There is a need to generate anti-CD3δ, anti-CD3ζ, anti-TCRα, and anti-TCRβ antibodies specific to Jamaican fruit bats to allow for the enhanced and accurate detection, visualization, and characterization of T lymphocytes and other CD3^+^ cell populations. The CD4 and CD8 bat specific antibodies generated in this study can be used to identify Th and CTL subsets and further elucidate the CD3 dimer stoichiometry in these populations of T cells.

Validation of cellular markers and reagents is a crucial first step in the investigative process that allows for the generation of informed conclusions, in any study, and hold a higher level of immanent need and accuracy in underdeveloped animal models. This same antibody validation methodology can be used to validated other cross-reactive antibodies across various species of bats. Additionally, this methodology can be applied to generate monoclonal antibodies to other targets, which is critical for bat researchers to rigorously validate and test cross-reactive antibodies to advance the field of bat immunology. The identification of novel CD3^+^ CD19^+^ T cells in the gastrointestinal tract and spleen of Jamaican fruit bats warrants further scrutiny to determine the physiological function and compounding affects that these cells may impose on the immune responses of Jamaican fruit bats. A CD19-expressing T cell population that activates or is aided in activation by the complement system could offer an explanation as to why bats may serve as reservoirs for viruses that cause high morbidity and mortality in humans and other mammalian populations because it could be a novel selection pressure for the evolution of bat-borne viruses.

## Materials and Methods

### Animals

All animal work was approved by the Colorado State University Institutional Animal Care and Use Committee. Mice and bats were handled in compliance with PHS policy and *Guide for the Care and Use of Laboratory Animals*. Mice were housed in microisolator cages (2 per cage), with a 12-hour light/dark cycle with access to food and water *ad libitum*. Jamaican fruit bats were housed in a free-flight captive breeding colony room with 12-hour light/dark cycle and provided fresh fruit (cantaloupe, watermelon, honeydew, banana) sprinkled with protein supplement each day.

### Pre-screening antibody cross-reactivity

Proteins that are more conserved between mammals are most likely to have cross-reactive mAb that are already commercially available. To identify such proteins, mouse and human protein sequences were BLASTed against the Jamaican fruit bat proteome. Alignment scores ≥50-80 were chosen as candidates to screen for cross-reactive mAb to Jamaican fruit bat splenocytes using flow cytometry. To further assess protein homology, protein sequences were downloaded into Geneious Prime and Geneious Protein Alignment BLOSUM62 and analysis was performed for human, mouse (*Mus musculus*), Jamaican fruit bat, common vampire bat (*Desmodus rotundus*), Chinese horseshoe bat (*Rhinolophus sinicus*), greater horseshoe bat (*Rhinolophus ferrumequinum*), Egyptian fruit bat (*Rousettus aegyptiacus*), Indian flying fox (*Pteropus medius*), and black flying fox (*Pteropus alecto*) for all CD3ε, CD3γ and CD19 protein isoforms (**Figure S1**) [29, 30]. BLOSUM62 homology precents were then averaged and protein alignment trees were generated. Greater than 60% protein homology determined if commercially available mouse and human antibody clones should be screened for cross-reactivity via flow cytometry. Additionally, protein sequences were uploaded to Phyre2 server to generate predicted 3D models of each protein [31]. The PDB files were then imported into Geneious Prime for BLOSUM62 alignment to identify identical protein alignment regions. These identical sites were colorimetrically annotated for extracellular regions (red) and transmembrane and intracellular regions (cyan).

### Generation of monoclonal antibodies

Jamaican fruit bat CD3γ, CD4, and CD8α protein sequences were submitted to the Phyre2 server to generate predicted 3D models of each protein. PDB files were imported into Geneious Prime, and the solvent-accessible peptide sequences were identified. Peptide sequences chosen for CD3γ (LKDQNIKWFKDGKEIMTNGNTWKLG), CD4 (ENRKVSVVKTRQDRR), and CD8α (PVFLMYIGTRTKIAEGLESQISGQKFQNDG) were synthesized with N-terminal Cys and conjugated to keyhole limpet hemocyanin (KLH; GenScript). Hybridomas were produced by immunizing two BALB/c mice for each target intraperitoneally with 25 μg of the peptide-KLH or IgG (H&L chains) emulsified in incomplete Freud’s adjuvant (IFA). Mice were identically boosted at 4 and 8 weeks after initial immunization. Twelve weeks post-initial immunization, mice were boosted with peptide in PBS and were euthanized 4 days later. Spleens were harvested and processed into a single cell suspensions and depleted of RBC with ammonium chloride solution (ACK Lysing buffer, Gibco). Splenocytes were then fused to Sp2/10-Ag14 cells with polyethylene glycol using the ClonaCell Hybridoma kit (StemCell Technologies). Fused cells were cultured overnight in T-75 flasks to recover, then mixed in methylcellulose containing HAT medium and plated onto five 100 mm Petri dishes for each hybridoma preparation for concurrent cloning and selection. Plates were incubated undisturbed for 14 days and colonies picked with sterile micropipettor tips and transferred to 96 well plates containing growth medium (StemCell Technologies). To screen hybridomas for mAb production, freshly harvested Jamaican fruit bat splenocytes depleted of RBC were incubated with hybridoma clone supernatants for 1 hr on ice. Splenocytes were subsequently washed and then stained with a secondary anti-mouse IgG antibody conjugated to FITC or Alexa Fluor 488 and analyzed by flow cytometry for reactive mAb.

### Tissue Extraction and Fixation

Adult male and female Jamaican fruit bats were placed under continual isoflurane anesthesia for perfusion. Intra-cardiac puncture was performed followed by intra-cardiac perfusion of the whole animal with 60 ml of perfusion buffer (1X PBS, 5 mM EDTA, 10 U/ml heparin sulfate). Tissue extraction was rapidly performed after perfusion with subsequent tissue fixation in 4% paraformaldehyde.

### Tissue Preparation and Automated High-throughput Immunofluorescence Staining

Paraffin embedded intestinal tissue was sectioned at 5 µm thickness and mounted onto polyionic slides. Sections were deparaffinized and labeled with immunofluorescence mAb on a fully automated Leica Bond RX_m_ robotic staining system. Epitope retrieval was performed through application of Bond Epitope Retrieval Buffer 1 (ER1) for 20 min at 95°C. Sections were then incubated with mAb directed toward epitopes of interest that were diluted in 0.1% Triton X100 in phosphate buffered saline (PBS). Antibodies were optimized for fluorescence staining at the following dilutions: CD3γ X-E2 conjugated to AF647 (1:50), CD3ε Hit3a conjugated to FITC (1:100), CD19 1D3 conjugated to PE-eFluor610 (1:500). Sections were stained with DAPI (Sigma) and mounted on glass coverslips with Prolong Gold anti-fade mounting media. Mounting medium was allowed to cure at room temperature in the dark for 24 hours. Sections were stored at 4°C until imaged.

### Detection and quantification of resident tissue immune cells

Full section montage images were acquired on a motorized stage scanning BX63 Olympus microscope equipped with a Hamamatsu ORCA-flash 4.0 LT CCD camera and collected using Olympus CellSens software (V3.1.1). Quantitative analysis was performed on tri-immunofluorescence labeled tissue section montages generated by compiling 100X and 200X images acquired using Olympus X-Apochromat 10X (0.40 N.A.) and 20X (0.8 N.A.) air objectives. Anatomical determination was performed by manual application of regions of interest (ROIs) based on tissue architecture and immunofluorescence labeled cells. High magnification inset images were acquired using an Olympus X-Apochromat 100X oil immersion objective (1.4 N.A.). Quantification of cell-specific populations within tissue sections was determined with the Count and Measure module of Olympus CellSens platform.

### Cell Sorting

Two male and two female bats were euthanized and spleens were removed and processed into a single cell suspensions. Splenocytes were then stained with Tonbo Ghost Violet 450 viability dye followed by staining with CD3γ (Clone: XE2:Alexa Fluor 647), CD3ε (Hit3a, FITC), and CD19 (1D3, PE-eFluor610). Cells were then sorted using a FACS Aria-III. UltraComp eBeads^TM^ Plus Compensation Beads were used for CD3ε, CD19; ArC^TM^ Amine Reactive Compensation Beads was used for Tonbo Ghost Violet 450; splenocytes for CD3γ were used for single color controls. FACSDiva Version 6.1.3 was used to calculated compensation. Gates were drawn using fluorescence minus one controls (FMOs). Cells were sorted into six categories by gating strategy: Not sorted (whole spleen), CD19^+^, CD3ε^+^, CD3γ^+^, and CD3ε^+^ CD3γ^+^. Sorted cells were then pelleted and supernatant was decanted. RNA was then extracted from the cell pellets using Trizol. RNA yields were determined by NanoDrop UV spectrometry (**Table 1** and compliance document available through the Flow Repository) [32, 33].

### Post-Hoc Cell Sorting Analysis of CD19^+^ CD3^+^ Cells

FCS files generated from cell sorting were analyzed in FlowJo 10.8.1. Single color controls as descried above were used to calculate compensation matrix in FlowJo. FCS files from all four bats were concatenated to better visualize CD19^+^ CD3^+^ cell populations. Gates were drawn using FMOs. Initial gating was: Intact > Single Cells > Live. CD19^+^ CD3^+^ cells were then viewed two ways: 1) first gating on CD3^+^ cells followed by CD19+ cells 2) first gating on CD19^+^ cells followed by CD3^+^ cells.

### Validation of Antibodies by RT-qPCR

Primer sets were designed for amplicons targeting β−actin, RPS18, CD3γ, CD3ε, CD4, CD8α, CD19, CD28, and CD45 (supplemental Table 1). Tonbo Low ROX One-step RT-qPCR kit was used. Primers were validated using unsorted splenocyte single cell suspension RNA, and the limit of quantification (LOQ) was determined for each primer set. CD3ε had the lowest LOQ, 11.222 ng of RNA. As such, 11.25 ng was the amount of RNA used for the RT-qPCR assay. RT-qPCR was performed on each sorted sample using all primer sets in duplicate. β-actin and RPS18 were used as reference genes. ΔΔCt method was then performed to calculate gene fold change and data was then normalized to whole spleen control. Two statistical tests were performed: one-way repeated measures ANOVA for pooled data output, and a two-way ANOVA to determine differences by sex (GraphPad Prism).

## Contributions

**Bradly E. Burke:** conceptualization, project design, performed experimentation, investigation, data curation, formal analysis, writing original draft, editing. **Savannah M. Rocha:** project design, performed experimentation, data curation, formal analysis, editing. **Miles Eckley:** performed experimentation, data curation, editing. **Corey Campbell:** performed experimentation, data curation, editing. **Elizabeth Creissen:** performed experimentation, data curation, editing. **Ronald B. Tjalkens:** editing, resources, supervision. **Marcela Henao-Tamayo:** project design, editing, resources, supervision. **Tony Schountz:** conceptualization, project design, investigation, editing, resources, supervision.

## Acknowledgements

We thank Colorado State University’s laboratory animal resources for animal support and Colorado State University’s Flow Cytometry and Cell Sorting Core facility for assistance (RRID# SCR_0220000_).

## Data Availability Statement

All flow cytometry data are available through the Flow Repository (https://flowrepository.org) with detailed methodology. For all other data please contact corresponding author Tony Schountz.

## Supplemental Figure Legends

**Supplemental Figure 1.**
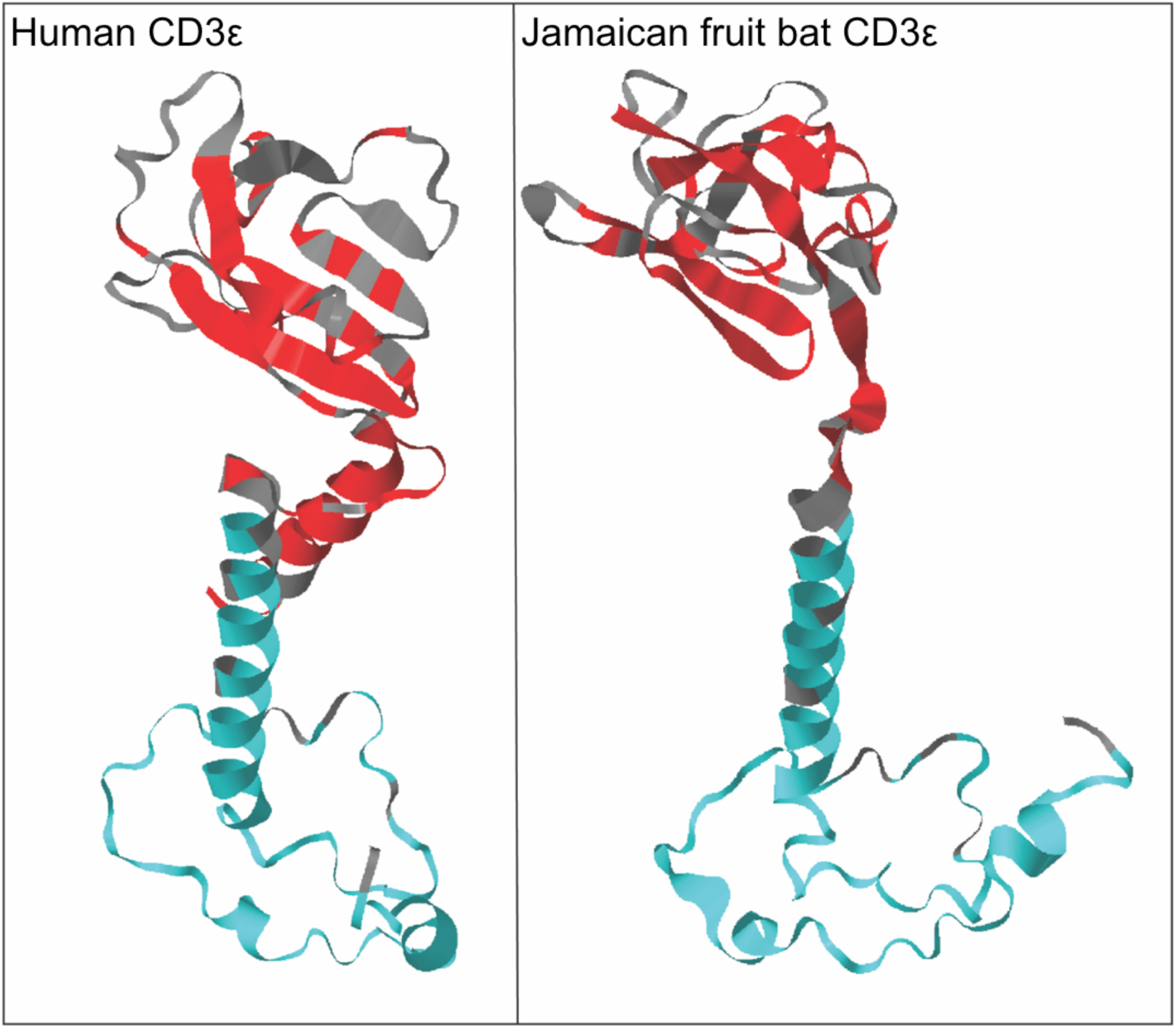
Three-dimensional modeling of human and Jamaican fruit bat CD3ε. Human CD3ε (NCBI accession NP_000724) and Jamaican fruit bat CD3ε (NCBI accession XP_037003230) amino acids sequences were processes through Phyre2 to generate predicted 3D structures. BLOSUM62 analysis between the two proteins indicated 75.5% homology with 133 (63.9%) identical sites, and colorimetric annotations highlight identical extracellular (red) and transmembrane and cytoplasmic (cyan) sites.

**Supplemental Figure 2.**
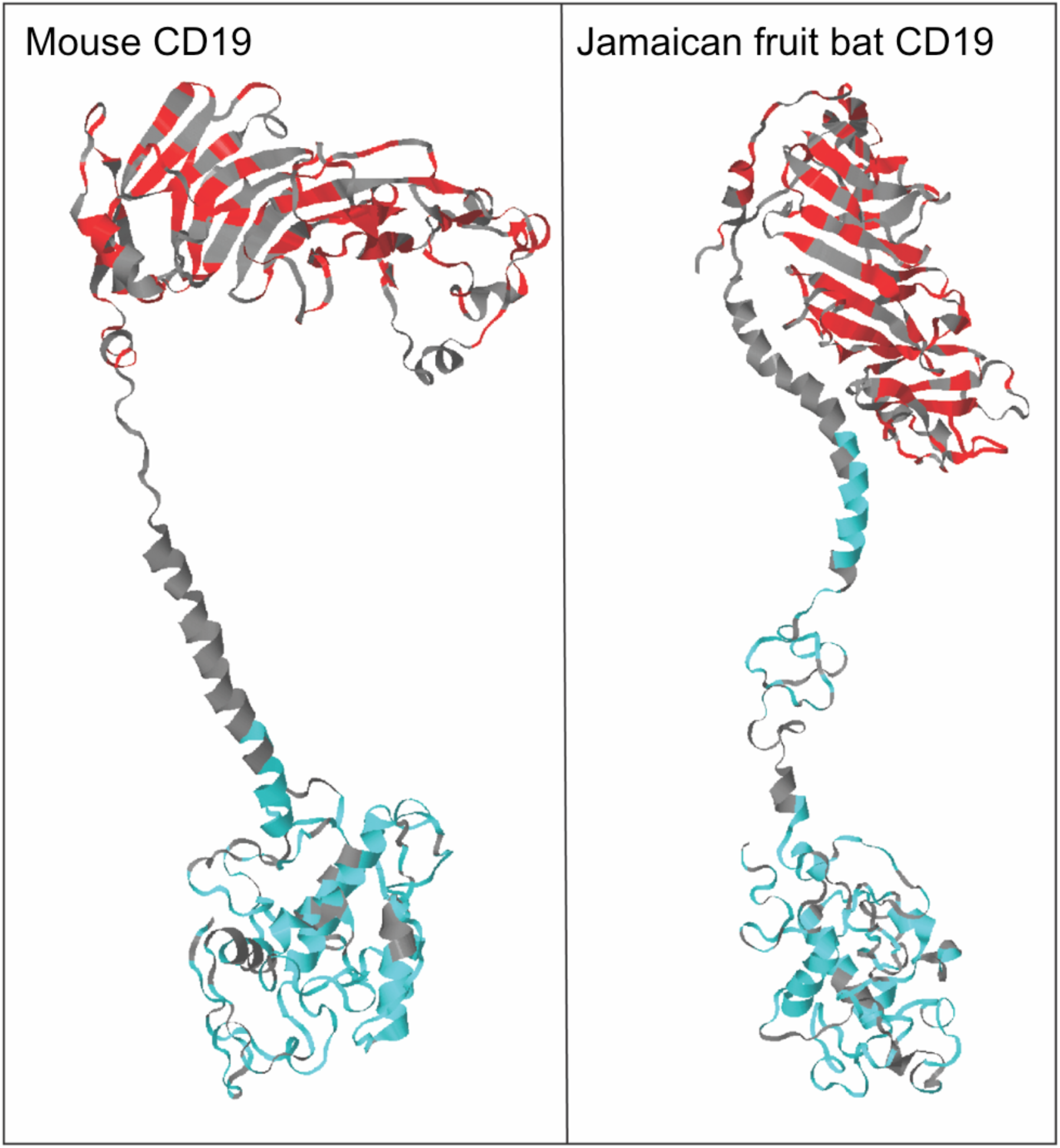
Three-dimensional modeling of murine and Jamaican fruit bat CD19. Mouse CD19 (NCBI accession AAA37390) and Jamaican fruit bat CD19 (NCBI accession XP_036995935) amino acids sequences were processes through Phyre2 intensive to generate predicted 3D structures. BLOSUM62 analysis between the two proteins indicated 67.1% homology with 336 (56.9%) identical sites, and colorimetric annotations highlight identical extracellular (red) and transmembrane and cytoplasmic (cyan) sites.

**Supplemental Figure 3.**
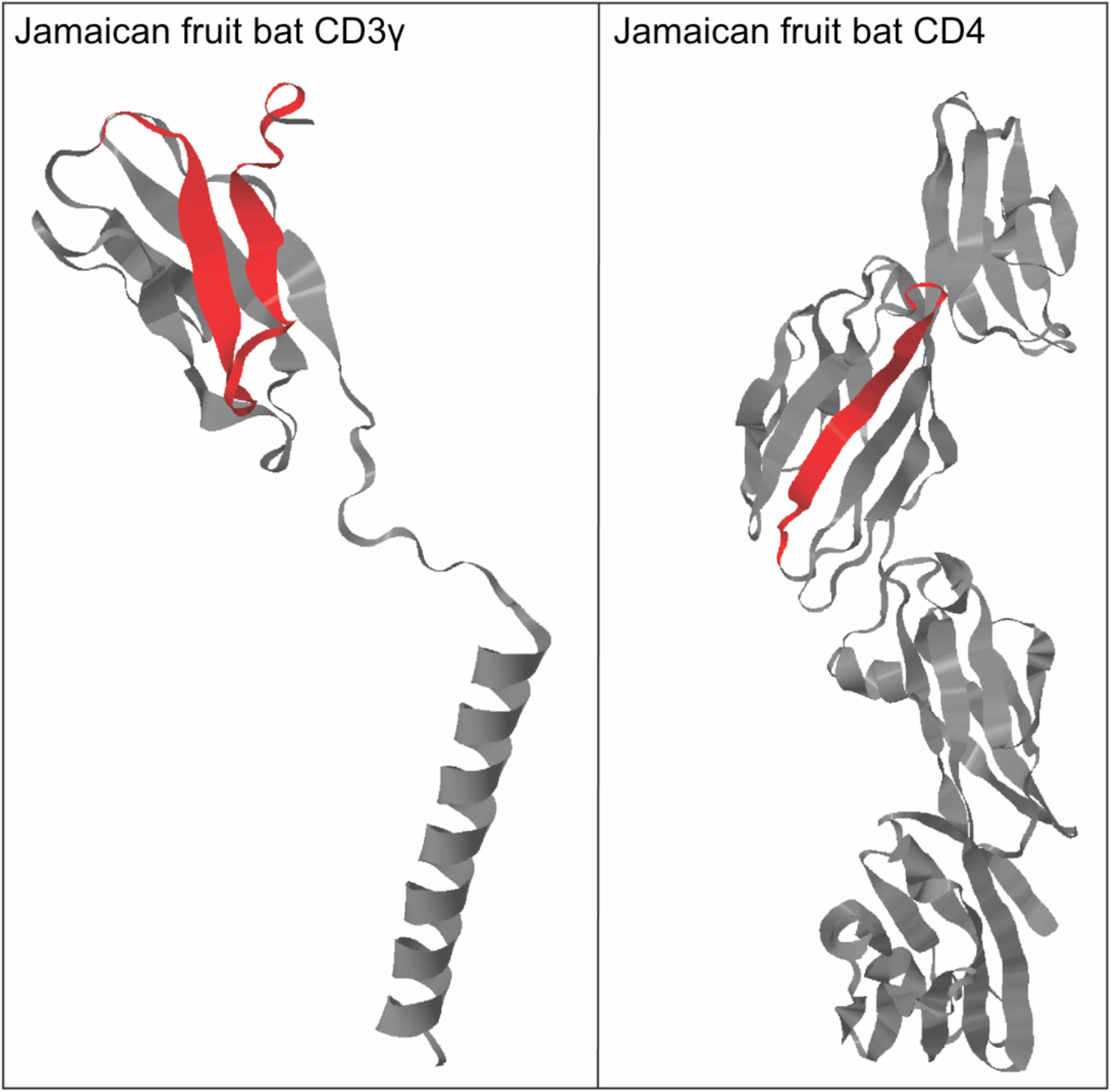
Three-dimensional modeling of murine and Jamaican fruit bat CD3γ and CD4. Jamaican fruit bat CD3γ (NCBI accession XP_037003233) and Jamaican fruit bat CD4 (NCBI accession XP_037016441) amino acids sequences were processes through Phyre2 intensive to generate predicted 3D structures. Red indicates solvent-accessible peptides that were used to generate KLH-peptide antigens for mouse immunizations to generate hybridomas.

**Supplemental Figure 4. Immunization dosing strategy and hybridoma production and screening.**

Two mice were immunized with KLH-conjugated CD3γ peptide and two mice were vaccinated with KLH-conjugated CD4 peptide in incomplete Freud’s adjuvant. Each peptide had a Cys residue added to the N-termini for KLH conjugation.

---

**Supplemental Table 1.**
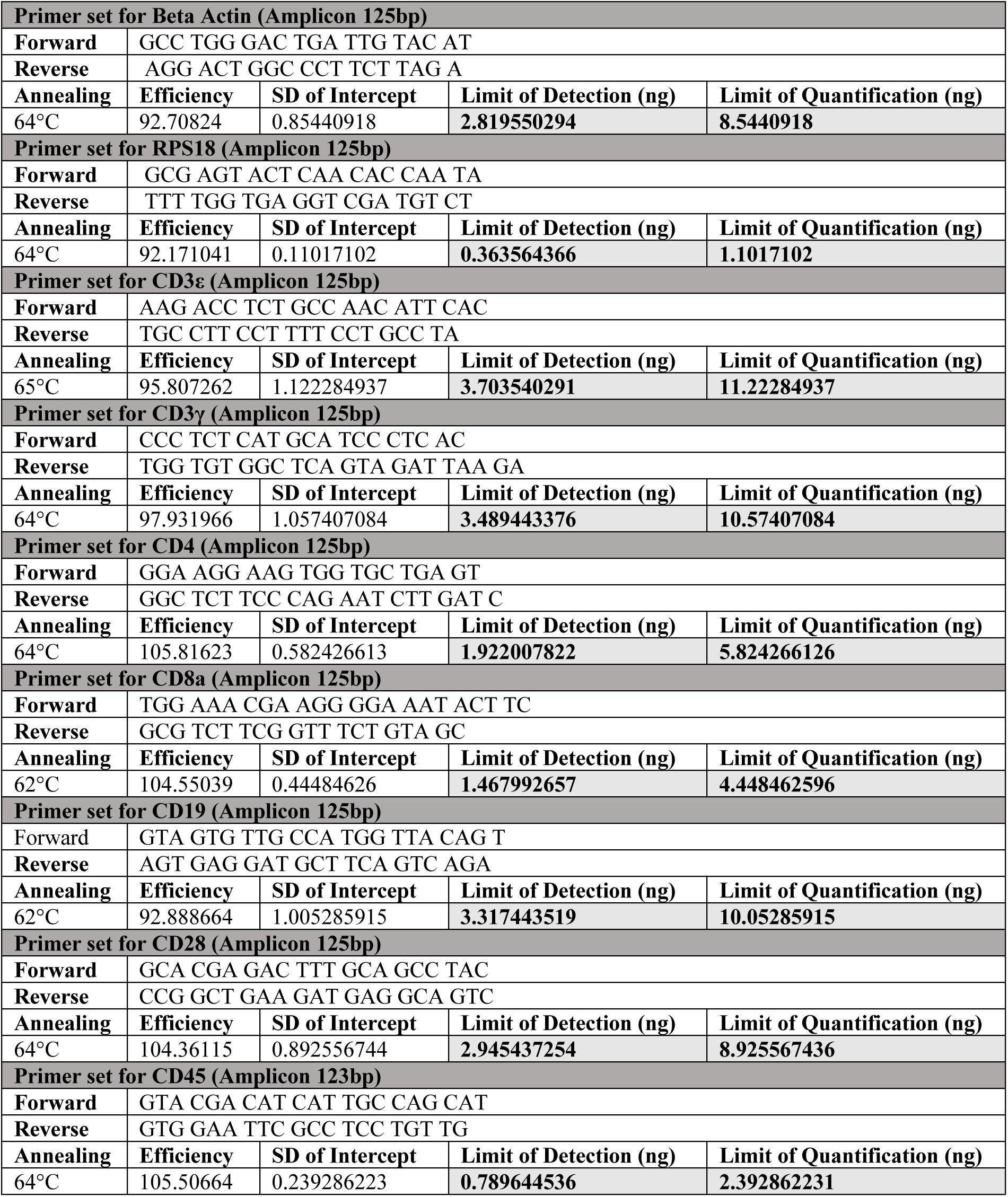

## References

1. Kunz, T.H., et al., Ecosystem services provided by bats, in Year in Ecology and Conservation Biology, R.S. Ostfeld and W.H. Schlesinger, Editors. 2011, Blackwell Science Publ: Oxford. p. 1–38.

2. Calisher, C.H., et al., Bats: Important reservoir hosts of emerging viruses. Clinical Microbiology Reviews, 2006. 19(3): p. 531-+.

3. Andersen, K.G., et al., The proximal origin of SARS-CoV-2. Nature Medicine, 2020. 26(4): p. 450–452.

4. Cheng, T.L., et al., The scope and severity of white-nose syndrome on hibernating bats in North America. Conservation Biology, 2021. 35(5): p. 1586–1597.

5. Zhou, P., et al., Contraction of the type I IFN locus and unusual constitutive expression of IFN-α in bats. Proc Natl Acad Sci U S A, 2016. 113(10): p. 2696–701.

6. Schountz, T., et al., Immunological Control of Viral Infections in Bats and the Emergence of Viruses Highly Pathogenic to Humans. Front Immunol, 2017. 8.

7. Bratsch, S., et al., The little brown bat, M. lucifugus, displays a highly diverse V-H, D-H and J(H) repertoire but little evidence of somatic hypermutation. Developmental and Comparative Immunology, 2011. 35(4): p. 421–430.

8. Baker, M., M. Tachedjian, and L. Wang, [Immunoglobulin heavy chain diversity in Pteropid bats: evidence for a diverse and highly specific antigen binding repertoire], in IMMUNOGENETICS. 2010. p. 173–184.

9. Max, E. and S. Fugmann, Fundamental immunology. 2013.

10. Pavlovich, S.S., et al., The Egyptian Rousette Genome Reveals Unexpected Features of Bat Antiviral Immunity. Cell., 2018. 173(5): p. 1098–1110.e18.

11. Munster, V.J., et al., Replication and shedding of MERS-CoV in Jamaican fruit bats (Artibeus jamaicensis). Scientific Reports, 2016. 6.

12. Malmlov, A., et al., Experimental Zika virus infection of Jamaican fruit bats (Artibeus jamaicensis) and possible entry of virus into brain via activated microglial cells. Plos Neglected Tropical Diseases, 2019. 13(2).

13. Cogswell-Hawkinson, A., et al., Tacaribe Virus Causes Fatal Infection of An Ostensible Reservoir Host, the Jamaican Fruit Bat. Journal of Virology, 2012. 86(10): p. 5791–5799.

14. Burke, B., et al., Regulatory T Cell-like Response to SARS-CoV-2 in Jamaican Fruit Bats (Artibeus jamaicensis) Transduced with Human ACE2. 2023, Cold Spring Harbor Laboratory.

15. E Reid, A.C.J., Jessica, Experimental rabies virus infection in Artibeus jamaicensis bats with CVS-24 variants. Journal of Neurovirology, 2001. 7(6): p. 511–517.

16. Colovai, A.I., et al., Flow cytometric analysis of normal and reactive spleen. Modern Pathology, 2004. 17(8): p. 918–927.

17. Nemoto, S., et al., OMIP-031: Immunologic Checkpoint Expression on Murine Effector and Memory T-Cell Subsets. Cytometry Part A, 2016. 89A(5): p. 427–429.

18. Dusoswa, S.A., J. Verhoeff, and J.J. Garcia-Vallejo, OMIP-054: Broad Immune Phenotyping of Innate and Adaptive Leukocytes in the Brain, Spleen, and Bone Marrow of an Orthotopic Murine Glioblastoma Model by Mass Cytometry. Cytometry Part A, 2019. 95A(4): p. 422–426.

19. Natalini, A., et al., OMIP-079: Cell cycle of CD4(+) and CD8(+) naive/memory T cell subsets, and of Treg cells from mouse spleen. Cytometry Part A, 2021. 99(12): p. 1171–1175.

20. Punt, J.A., et al., STOICHIOMETRY OF THE T-CELL ANTIGEN RECEPTOR (TCR) COMPLEX - EACH TCR/CD3 COMPLEX CONTAINS ONE TCR-ALPHA, ONE TCR-BETA, AND 2 CD3-EPSILON CHAINS. Journal of Experimental Medicine, 1994. 180(2): p. 587–593.

21. Delahera, A., et al., STRUCTURE OF THE T-CELL ANTIGEN RECEPTOR (TCR) - 2 CD3-EPSILON SUBUNITS IN A FUNCTIONAL TCR/CD3 COMPLEX. Journal of Experimental Medicine, 1991. 173(1): p. 7–17.

22. Periasamy, P., et al., Studies on B Cells in the Fruit-Eating Black Flying Fox (Pteropus alecto). Frontiers in Immunology, 2019. 10.

23. Rizzo, K., et al., Novel CD19 Expression in a Peripheral T Cell Lymphoma: A Flow Cytometry Case Report with Morphologic Correlation. Cytometry Part B-Clinical Cytometry, 2009. 76B(2): p. 142–149.

24. Schultz, L., et al., Identification of dual positive CD19+/CD3+T cells in a leukapheresis product undergoing CAR transduction: a case report. Journal for Immunotherapy of Cancer, 2020. 8(2).

25. Bradford, C., R. Jennings, and J. Ramos-Vara, Gastrointestinal leiomyosarcoma in an Egyptian fruit bat (Rousettus aegyptiacus). Journal of Veterinary Diagnostic Investigation, 2010. 22(3): p. 462–465.

26. McLelland, D.J., C.J. Dutton, and I.K. Barker, Sarcomatoid carcinoma in the lung of an Egyptian fruit bat (Rousettus aegyptiacus). Journal of Veterinary Diagnostic Investigation, 2009. 21(1): p. 160–163.

27. Matsumoto, A.K., et al., Intersection of the complement and immune systems: a signal transduction complex of the B lymphocyte-containing complement receptor type 2 and CD19. The Journal of experimental medicine, 1991. 173(1): p. 55–64.

28. Carter, R.H. and D.T. Fearon, CD19 - LOWERING THE THRESHOLD FOR ANTIGEN RECEPTOR STIMULATION OF LYMPHOCYTES-B. Science, 1992. 256(5053): p. 105–107.

29. Geneious Prime 2022.2.1 (https://www.geneious.com).

30. Henikoff, S. and J.G. Henikoff, AMINO-ACID SUBSTITUTION MATRICES FROM PROTEIN BLOCKS. Proceedings of the National Academy of Sciences of the United States of America, 1992. 89(22): p. 10915–10919.

31. Kelley, L.A., et al., The Phyre2 web portal for protein modeling, prediction and analysis. Nature Protocols, 2015. 10(6): p. 845–858.

32. Ja, L., et al., MIFlowCyt: The Minimum Information About a Flow Cytometry Experiment. Cytometry. Part A : the journal of the International Society for Analytical Cytology, 2008. 73(10).

33. Spidlen, J., K. Breuer, and R. Brinkman, Preparing a Minimum Information about a Flow Cytometry Experiment (MIFlowCyt) compliant manuscript using the International Society for Advancement of Cytometry (ISAC) FCS file repository (FlowRepository.org). Current protocols in cytometry, 2012. **Chapter** 10: p. Unit 10.18–Unit 10.18.

